# Knowledge and attitudes among life scientists towards reproducibility within journal articles: a research survey

**DOI:** 10.1101/581033

**Authors:** Evanthia Kaimaklioti Samota, Robert P. Davey

## Abstract

We constructed a survey to understand how authors and scientists view the issues around reproducibility, focusing on interactive elements such as interactive figures embedded within online publications, as a solution for enabling the reproducibility of experiments. We report the views of 251 researchers, comprising authors who have published in *eLIFE* Sciences, and those who work at the Norwich Biosciences Institutes (NBI). The survey also outlines to what extent researchers are occupied with reproducing experiments themselves. Currently, there is an increasing range of tools that attempt to address the production of reproducible research by making code, data, and analyses available to the community for reuse. We wanted to collect information about attitudes around the consumer end of the spectrum, where life scientists interact with research outputs to interpret scientific results. Static plots and figures within articles are a central part of this interpretation, and therefore we asked respondents to consider various features for an interactive figure within a research article that would allow them to better understand and reproduce a published analysis. The majority (91%) of respondents reported that when authors describe their research methodology (methods and analyses) in detail, published research can become more reproducible. The respondents believe that having interactive figures in published papers is a beneficial element to themselves, the papers they read as well as to their readers. Whilst interactive figures are one potential solution for consuming the results of research more effectively to enable reproducibility, we also review the equally pressing technical and cultural demands on researchers that need to be addressed to achieve greater success in reproducibility in the life sciences.

## Introduction

Reproducibility is a defining principle of scientific research, and broadly refers to the ability of researchers, other than the original researchers, to achieve the same findings using the same data and analysis [1]. However, irreproducible experiments are common across all disciplines of life sciences [2]. A recent study showed that 88% of drug-discovery experiments could not be reproduced even by the original authors, in some cases forcing retraction of the original work [2]. Irreproducible genetic experiments with weak or wrong evidence can have negative implications for our healthcare [3]. For example, 27% of mutations linked to childhood genetic diseases cited in literature have later been discovered to be common polymorphisms or misannotations [4]. While irreproducibility is not confined to biology and medical sciences [5], irreproducible biomedical experiments pose a strong financial burden on society; an estimated $28 billion was spent on irreproducible biomedical science in 2015 in the USA alone [6].

Reproducibility should inevitably lead to robust science, relating to the way in which conclusions rely on specific analyses or procedures undertaken on experimental systems. Unfortunately, the community has yet to reach consensus on how we traverse the space of re-use, re-analysis and re-interpretation of scientific research to try to define suitable overarching definitions for reproducibility. Thus, there are different definitions of *reproducibility* used in the literature [7,8], some of which contradict one another. A recent exhaustive review has also documented this problem [9], so our survey and results do need to be contextualised somewhat by this lack of consensus. The terms *repeatability, replicability* and *reproducibility* are also occasionally confused [10,11], therefore it is important to differentiate these terms from each other.

1. **Repeatability** The original researchers using the same data, running precisely the same analysis and getting the same results, on multiple runs [8].
2. **Replicability** Different teams performing different experimental setups and using independent data, achieving the same result as the original researchers, on multiple trials [11-13].
3. **Reproducibility** Different teams re-running the same analysis with the same data and getting the same result [1,11-13].

It is argued that in many science disciplines replicability is more desirable than reproducibility because a result needs to be corroborated independently before it can be generally accepted by the scientific community [12,14]. However, reproducibility can serve as a cost-effective way of verifying results prior to replicating results [14].

*Computational reproducibility*, or reproducible computational research, refers to the reproducibility of computational experiments, where an independent team can produce the same result utilising the data and computational methods (code and workflow) provided by the original authors [9,13,15-17]. Computational reproducibility is influenced by both technical and cultural (social) factors [13,17,18]. Technical challenges to computational reproducibility include poorly written, incorrect, or unmaintained software, changes in software libraries on which tools are dependent, or incompatibility between older software and newer operating systems [19]. Cultural factors that challenge computational reproducibility include the attitudes and behaviours of authors when performing and reporting research. Examples include authors not providing sufficient descriptions of methods and being reluctant to publish original data and code under FAIR (Findable, Accessible, Interoperable, and Reusable) principles[13,20,21]. Other cultural factors include favouring of high prestige or high impact science publications over performing rigorous and reproducible science (which tends to be improved by open access policies) [22,23]. We refer to the cultural factors affecting computational reproducibility as the *culture of reproducibility* [12].

Several projects have attempted to address some of the technical aspects of reproducibility by making it easier for authors to disseminate fully reproducible workflows and data, and for readers to perform computations. For example: F1000 Living Figure [24] and re-executable publications [25-28] using Plotly (plot.ly) and Code Ocean widgets (codeocean.com); Whole Tale Project [29]; ReproZip project [30]; Python-compatible tools and widgets (interactive widgets for Jupyter Notebooks with Binder); Zenodo (zenodo.org) and FigShare (figshare.com) as examples of open access repositories for scientific content (including datasets, code, figures, reports); Galaxy [31]; CyVerse (formerly iPlant Collaborative) [32]; ^my^[Experiment](http://www.myexperiment.org/) [33]; UTOPIA [34,35]; GigaDB [36]; Taverna [37-39]; workflow description efforts such as the Common Workflow Language [40]; and Docker (docker.com), Singularity (singularity.lbl.gov) [41], and other container systems. Even though these tools are widely available, and seem to address many of the issues of *technical reproducibility* and the *culture of reproducibility*, they have not yet become a core part of the life sciences experimental and publication lifecycle. There is an apparent disconnection between the development of tools addressing reproducibility and their use by the wider scientific and publishing communities who might benefit from them.

This raises the question of “how do researchers view their role in the production and consumption of scientific outputs?” A common way for researchers to quickly provide information about their data, analysis and results is through a figure or graph. Scientific figures in publications are commonly presented as static images. Access to the data (including the raw, processed and/or aggregated data), analysis, code or description of how the software was used that produced the figure are not available within the static images [42-48]. This can be especially pertinent to figures that have thousands or millions of points of data to convey [42]. In order for readers to interrogate published results in more detail, examine the transparency and reproducibility of the data and research, they would need to download a complete copy of the data, code, and any associated analysis methodology (data pre-processing, filtering, cleaning, etc) and reproduce this locally, provided all those elements are available and accessible [49]. Computational analyses often require running particular software which might require configuration and parameterisation, as well as library dependencies and operating system prerequisites. This is a time-consuming task and achieving reproducibility of computational experiments is not always possible [49,50]. Thus, solutions that automatically reproduce computational analyses and allow the investigation of the data and code presented in the figure in detail would be advantageous [12,26,42].

Many solutions now exist that allow for the reproducibility of computational analyses outside the research paper, and are typically supplied as links within the research paper or journal website redirecting to many different types of computational systems, such as Galaxy workflows, Binder interactive workspaces converted by GitHub repositories with Jupyter notebooks [51], and myExperiment links [33]. The endpoint of these analyses are often graphical figures or plots, and these may well be interactive, thus allowing modification of plot type, axes, data filtering, regression lines, etc. Whilst these figures may well be interactive in that a user can modify some part of the visualisation, this does not implicitly make the data or code that produced that figure more available, and hence more reproducible.

Technologies that can expose code, data and interactive figures are now mature. For example, Jupyter notebooks are built up of executable “cells” of code which can encapsulate a link to a data file hosted on a cloud service, code to get and analyse this data file, and then produce an interactive figure to interpret the dataset. Again, this is somewhat disconnected from the actual research publication. However, as technology has progressed in terms of available storage for data, computational power on the web through cloud services, and the ability of these services to run research code, we are now coming to the point where the production of interactive figures within publications themselves is achievable. These interactive figures which would inherently have access to the underlying data and analytical process can provide users with unique functionality that can help increase the reproducible nature of the research. This combination of code, data, analysis, visualisation and paper are examples of “executable documents” [52,53].

Interactive figures within executable documents therefore have incorporated data, code and graphics so that when the user interacts with the figure, perhaps by selecting a cluster of data points within a graph, the user could then be presented with the data that underlies those data points. Similarly, a user could make changes to the underlying parameters of the analysis, for example modifying a filter threshold, which would ultimately make changes to the visualisation of the figure or the document itself [42-48]. By means of an example, an executable document could represent an interactive figure showing a heat map of gene expression under different stress conditions. In a traditional article, the user would be tasked with finding references to the datasets and downloading them, and subsequently finding the code or methodology used to analyse the data, and retrace the original authors steps (if the code and data were available at all). Within an interactive figure in an executable document, a user could select a particular gene of interest by clicking on the heatmap, and viewing the gene expression information within a pop-up browser window. Whilst this is useful for general interpretation, to achieve reproducibility this pop-up window would provide a button that allows the user to pull the sequencing read data that was the basis for the results into a computational system in order to re-run the differential expression analysis. This raises many questions around how this infrastructure is provided, what technologies would be used to package up all elements needed for reproducibility, the subsequent costs of running the analysis, and so on.

These caveats aside, interactive figures within executable documents can benefit the reader for the consumption of the research outputs in an interactive way, with easy access to the data and removing the need for installing and configuring code and parameters for reproducing the computational experiments presented in the figure within the publication [26]. The aforementioned solutions would not only be helpful to the readers of papers [54], but benefit the peer review process [42].

There have been efforts to make the connection between production and consumption of research outputs within online publications. One of the first interactive figures to have been published in a scholarly life sciences journal is the *Living Figure* by Björn Brembs and Julien Colomb which allowed readers to change parameters of a statistical computation underlying a figure [25]. *F1000Research* has now published more papers that include Plotly graphs and Code Ocean widgets in order to provide interactivity and data and code reproducibility from within the article figures [28,52]. The first prototype of *eLIFE’s* computationally reproducible article aims to convert manuscripts created in a specific format (using the Stencila Desktop, stenci.la, and saved as a Document Archive file) into interactive documents allowing the reader to “play” with the article and its figures when viewed in a web browser [53]. The Manifold platform (manifoldapp.org) allows researchers to show their research objects alongside their publication in an electronic reader, whilst including some dynamic elements. The *Cell* journal included interactive figures in a paper using Juicebox.js for 3D visualisation of Hi-C data (http://aidenlab.org/juicebox/) [55,56].

Whilst there are few incentives to promote the culture of reproducibility [57,58], efforts in most science domains are being made to establish a culture where there is an expectation to share data for all publications according to the FAIR principles. The implementation of these principles is grounded in the assumption that better reproducibility will benefit the scientific community and the general public [59,60]. Studies have suggested that reproducibility in science is a serious issue with costly repercussions to science and the public [61,62]. Whilst there have been survey studies canvassing the attitudes of researchers around reproducibility in other disciplines to some extent [20,63,64], fewer studies have investigated the attitudes and knowledge of researchers around reproducibility in the life sciences [20]. In particular, minimal research has been conducted into the frequency of difficulties experienced with reproducibility, the perception of its importance, and preferences with respect to potential solutions among the life sciences community.

This paper presents a survey that was designed to assess researcher understanding of the concepts of reproducibility and to inform future efforts in one specific area: to help researchers be able to reproduce research outputs in publications. The development of tools, one example of which are interactive figures within journal publications, may better meet the needs of producers and consumers of life science research. Our survey is limited in that we do not assess how open access tools for the production of reproducible research outputs compare, but how the consumption of research information through interactive means is regarded. We constructed the survey in order to understand how the following are experienced by the respondents:

- *Technical factors affecting computational reproducibility*: issues with accessing data, code and methodology parameters, and how solutions such as interactive figures could promote reproducibility from within an article.
- *Culture of reproducibility*: attitudes towards reproducibility, the social factors hindering reproducibility, and interest in how research outputs can be consumed via interactive figures and their feature preferences.

## Methods

### Population and sample

The data were analysed anonymously, nonetheless, we sought ethical approval. The University of East Anglia Computing Sciences Research Ethics Committee approved this study (CMPREC/1819/R/13). Our sample populations were selected to include all life sciences communities across levels of seniority, discipline and level of experience with the issues we wished to survey. The first survey was conducted in November 2016 and sent out to 750 researchers working in the Norwich Biosciences Institutes (NBI) at a post-doctoral level or above. We chose to survey scientists of post-doctoral level or above, as these scientists are more likely to have had at least one interaction with publishing in scientific journals. The NBI is a partnership of four UK research institutions: the Earlham Institute (formerly known as The Genome Analysis Centre), the John Innes Centre, the Sainsbury Centre, and the Institute of Food Research (now Quadram Institute Bioscience). Invitations to participate were distributed via email, with a link to the survey. The second survey, similar to the first but with amendments and additions, was distributed in February 2017 to a random sample of 1651 researchers who had published papers in the *eLIFE* journal. Further information about the *eLIFE* sample is found in Supplementary section 3. Invitations to participate were sent using email by *eLIFE* staff. We achieved a 15% (n=112) response rate from the NBI researchers and an 8% response rate from the *eLIFE* survey (n=139). Table 1 shows the survey questions. Questions were designed to give qualitative and quantitative answers on technical and cultural aspects of reproducibility. Questions assessed the frequency in difficulties encountered in accessing data, the reasons for these difficulties, and how respondents currently obtain data underlying published articles. They measured understanding of what constitutes reproducibility of experiments, interactive figures, and computationally reproducible data. Finally, we evaluated the perceived benefit of interactive figures and of reproducing computational experiments, and which features of interactive figures would be most desirable.

**Table 1:** Questions used to survey the knowledge of respondents about research reproducibility.

|  | Survey questions |
| --- | --- |
| 1 | How often do you encounter difficulties with working with bioinformatic analysis tools (that are not your own)? (Problems such as: installing, configuring, running the software, working with command-line software)? |
| 2 | How difficult is it to source (or access) the data presented in published papers? |
| 3 | What difficulties have you encountered in accessing the data described in published papers? |
| 4 | How are you currently sourcing the data (if applicable)? Select all that apply to you. |
| 5* | What is your current understanding of reproducibility of experiments? Please select any that apply. Should you wish to add any additional information, please add it to the “Other” box. |
| 6* | Have you ever tried reproducing any published results? Please select the answer that applies best for you. |
| 7* | In your opinion, what could be done to make published research more reproducible? Other please specify (free text answer). |
| 8 | When thinking about *interactive figures*, what comes to your mind? (please describe what you understand of what an interactive figure to be, its features, and where you have seen such a |
|  | feature before if applicable). |
| 9 | An *interactive figure* is a figure within a paper that is dynamic and becomes “live” when the user interacts with it and where the data displayed changes according to various parameter options. Which of the following features of an interactive figure tool would be good to have? Please rank them in the order of preference, where 1 is the most preferred feature, and 11 the least preferred feature. |
| 10 | What other features an *interactive figure* could have that were not mentioned in the previous question? |
| 11 | Do you perceive a benefit in being able to publish *interactive figures*? |
| 12 | Does the provision or option of an *interactive figure* in the paper affect your decision in choosing the publishing journal or publisher? |
| 13 | Have you heard of the term *computationally reproducible data*, and do you understand what the term means? If answered yes or unsure, please explain what you understand from the term. |
| 14 | Would you benefit from being able to automatically reproduce computational experiments or other analyses (including statistical tests) described within a paper? |
| 15 | How often do you work with *bioinformatic analysis tools* (e.g. assemblers, aligners, structure modelling)? |
| 16 | Have you received any of the following training? Training whether formal or informal (training through a colleague etc.). |
| 17 | Which of the following type(s) of data do you work with? |

\*Questions indicated with an asterisk were only available to the *eLIFE* survey. Answer options to the questions are shown in Supplementary Section 1.

### Validation of the survey design

We undertook a two-step survey: firstly NBI, then *eLIFE* interactions leading to additional questions. We tested the initial survey on a small cohort of researchers local to the authors to determine question suitability and flow. We designed the surveys in accordance with the Standards for Reporting Qualitative Research (SRQR) [65]. Our study was not pre-registered, but we confirm that the reported analyses are the only analyses that were conducted and that the reported variables were the only variables that were measured.

### Statistical analysis

Results are typically presented as proportions of those responding, stratified by the respondent’s area of work, training received, and version of the survey as appropriate. Chi-square tests for independence were used to test for relationships between responses to specific questions, or whether responses varied between samples. The analysis was conducted using R (version 3.5.2; R Core Team, 2018) and Microsoft Excel. All supplementary figures and data are available on Figshare (see Data Availability).

We assessed if there was a significant difference in the ability and willingness to reproduce published results between the cohort of *eLIFE* respondents who understand the term “computationally reproducible data” and those who do not and whether training received, had an effect. Given the free-text responses within the “unsure” group as to the understanding of this term, where many understood the term, we did not include those that replied, “unsure” (see Section “Understanding of reproducibility, training and successful replication” below). The respondents who chose “yes tried reproducing results, but unsuccessfully”, “have not tried to reproduce results” and “it is not important to reproduce results” were grouped together under “unsuccessfully”.

## Results

### Characteristics of the sample

Figure 1 shows the distribution of areas of work of our respondents, stratified by survey sample. Genomics (proportion in the whole sample = 22%), biochemistry (17%), and computational biology (15%) were the most common subject areas endorsed in both NBI and *eLIFE* samples. With regard to how often respondents use bioinformatics tools, 25% replied “never”, 39% “rarely”, and 36% “often”. Many (43%) received statistical training, (31%) bioinformatic training, (20%) computer science training.

**Figure 1:**
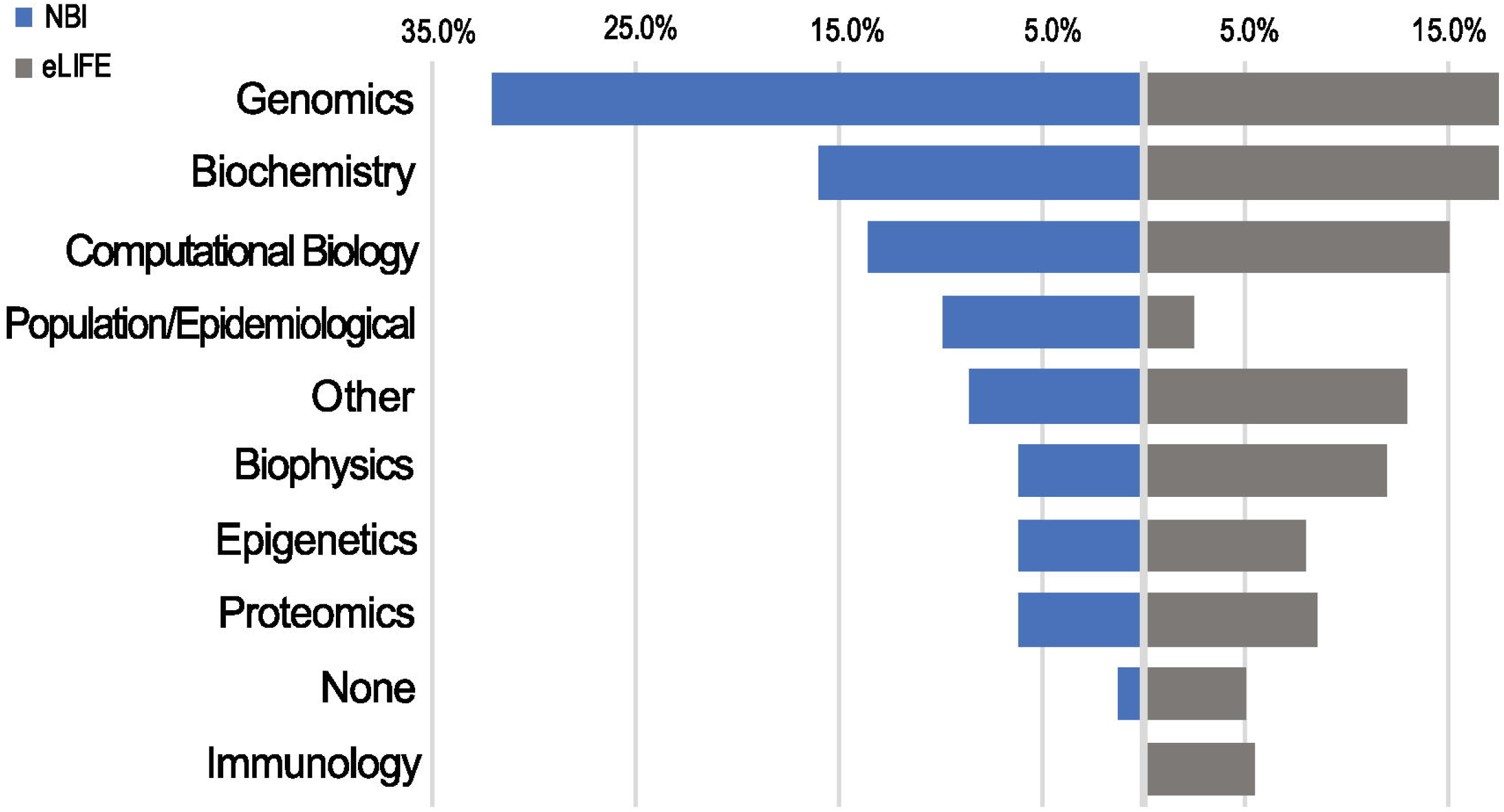
Data types used by NBI and *eLIFE* respondents. Responses were not mutually exclusive. Data type choices were the same as the article data types available in the *eLIFE* article categorisation system.

### Access to data and bioinformatics tools

In both samples, 90% of those who responded, reported having tried to access data underlying a published research article (Fig. 2). Of those who had tried, few had found this “easy” (14%) or “very easy” (2%) with 41% reporting that the process was “difficult” and 5% “very difficult”. Reasons for difficulty were chiefly cultural (Fig. 2), in that the data was not made available alongside the publication (found by 75% of those who had tried to access data), or authors could not be contacted or did not respond to data requests (52%). Relatively few found data unavailable for technical reasons of data size (21%), commercial sensitivity (13%) or confidentiality (12%). With respect to data sources, 57% of the total sample have used open public databases, 48% reported data was available with a link in the paper, and 47% had needed to contact authors.

**Figure 2.**
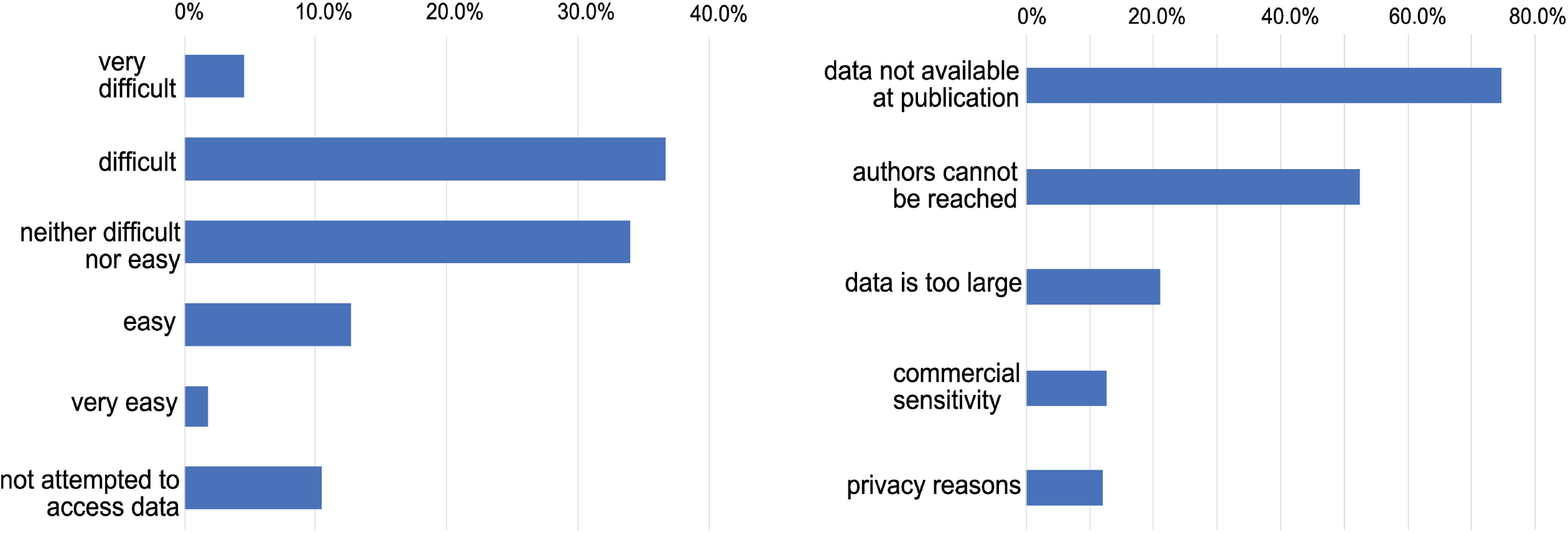
Left panel: Difficulty encountered accessing data underlying published research. Whether respondents have attempted to access data underlying previous publications and the level of difficulty typically encountered in doing so. **Right panel: Reasons given for difficulty accessing data**. The reasons given by respondents for being unable to access data (restricted to those who have attempted to access data).

Very few of the respondents either “never” (2%) or “rarely” (8%) had problems with running, installing, configuring bioinformatics software. Problems with software were encountered “often” (29%) or “very often” (15%) suggesting that nearly half of respondents regularly encountered technical barriers to computational reproducibility.

### Understanding of reproducibility, training and successful replication

The majority of respondents reported that they understood the term “reproducibility of experiments” and selected the explanation for the term as defined in the introduction above, which corresponds to the most established definitions of reproducibility [11-13]. It is important to state that for this question, we allowed for respondents to choose more than one answer, as we recognise the limitation that there is no standard and accepted definition for reproducibility, as well as the familiarity of the term between scientists from different backgrounds can differ. The first three definitions are plausible definitions for reproducibility. Given the results, we can assume that some of the respondents chose both correct and wrong definitions. The majority of the answers (77%) included the definition of reproducibility as we define it in the manuscript. However, by looking into the individual responses (n=54), 11.1% (n=6) of respondents chose only option A thus appeared to understand that this matched the definition of reproducibility, as we state in the manuscript. 5.5% (n=3) chose only option D, which is incorrect. The majority of people (57%, n=23) picked any of A, B, or C and did *not* pick D, which seems to suggest that they understand that replicability is not reproducibility, but they are still not clear on exact definitions, which matches the general lack of consensus [7,8,10]. Just over a third (37%, n=20) picked one or all of A, B and C, *and* picked D, which seems to suggest that they didn’t understand the difference between reproducibility and replicability at all and considered any form of repeating a process could be classed as “reproducibility of experiments (see Supplementary table 4).

Most (52%) participants provided a different interpretation of the term “computationally reproducible data” to our interpretation, while 26% did know and 22% were unsure. We received several explanations (free text responses) of the term of which the majority were accurate (Supplementary section 2, free responses to question 13). We assign meaning to the term as data as an output (result) in a computational context, which was generated when reproducing computational experiments. Although the term “computationally reproducible data” is not officially defined, other sources and studies have referred to the concept of data that contributes to computational reproducibility [26,66-70]. From the unsure responses (n=30), we categorised those that gave free-text responses (70%, n=21, see Supplementary section 2, free responses) into whether they did actually understand the term, those that did not understand the term, and those that did not give any free text. The majority of respondents that chose “unsure” and gave a free text response (71%, n=15) did understand the term “computationally reproducible data”. The remaining 29% (n=6) did not understand the term correctly.

Some (18%) reported not attempting to reproduce published research. Very few (n = 5; 6%) of the sample endorsed the option that “it is not important to reproduce other people’s published results” (Supplementary figure 1). Even though the majority (60%) reported successfully reproducing published results, almost a quarter of the respondents found that their efforts to reproduce any results were unsuccessful (23%). Table 2 shows the ability of respondents in reproducing experiments stratified by the understanding of the term “computationally reproducible data” and the training received (bioinformatics, computer science, statistics). We found a significant difference between the ability to reproduce published experiments and knowing the meaning of the term “computationally reproducible data”. Among the 25 respondents who understood the term “computationally reproducible data”, 18 (72%) had successfully reproduced previous work, compared to only 26 (52%) of the 50 who responded that they did not understand the term (Chi-square test for independence, p = 0.048). Taking their training background into account did not show any significant difference. However, when testing with the responses “yes tried reproducing results, but unsuccessfully”, “have not tried to reproduce results” and “it is not important to reproduce results” (not grouped together under “unsuccessfully” in order to get an indication of how willingness and success together differed between the training groups), we found a significant difference (see Supplementary Table 1). The distribution of the training variable with those who received computer science training and those without was significantly different (Fisher exact test for independence, p = 0.018). It appears that respondents with computer science training are less likely to have tried to reproduce an experiment but be more likely to succeed when they did try.

**Table 2:** Success in reproducing any published results stratified by their knowledge of the term “computationally reproducible data” and training received.

|  | Number<br>(% of the<br>total sample) | Success in reproducing any published results |  |  |
| --- | --- | --- | --- | --- |
| Variable |  | Successful (%<br>within<br>variable) | Not Successful*<br>(% within<br>variable) | P-value |
| **Knowledge of the term**<br>**“computationally reproducible**<br>**data” ($n = 75$)** |  |  |  |  |
| Yes | 25 (33.3) | 18 (72) | 7 (28) | 0.048** |
| No | 50 (66.7) | 24 (48) | 26 (33) |  |
| **Training ($n = 90$)** |  |  |  |  |
| Bioinformatics | 42 (46.7) | 26 (61.9) | 16 (38.1) | 0.73 |
| *Not trained in Bioinformatics* | 48 (53.3) | 28 (58.3) | 20 (41.7) |  |
| Computer Science | 33 (36.7) | 21 (63.6) | 12 (36.4) | 0.59 |
| *Not trained in Computer Science* | 57 (63.3) | 33 (57.9) | 24 (42.1) |  |
| Statistics | 71 (78.9) | 42 (59.2) | 29 (40.8) | 0.75 |
| *Not trained in Statistics* | 19 (21.1) | 12 (63.2) | 7 (36.8) |  |
| No training | 10 (11.1) | 6 (60) | 4 (40) | 0.73*** |
| *All other training* | 80 (88.8) | 48 (60) | 32 (40) |  |

*n* is different for the two variables as not all participants answered all the questions
\*Unsuccessful includes answers: “Yes, I have tried reproducing published results, but I have been unsuccessful in producing any results, or the same results”, “No, I have never tried reproducing any published results” and “It is not important to reproduce other people’s published results”
\*\*Statistically significant at the level of $P < 0.05$
\*\*\*Chi-square statistic with Yates correction, applied when expected frequencies were lower than 5

There was no evidence for a difference in the ability and willingness to reproduce published results between the respondents who use bioinformatics tools often, and those who use them rarely or never (Supplementary Table 2). The majority of the respondents who use bioinformatics tools often were coming from the scientific backgrounds of Biophysics, Biochemistry, Computational Biology and Genomics. Most of the respondents who answered “reproducibility is not important” and “haven’t tried reproducing experiments” were scientists coming from disciplines using computational or bioinformatics tools “rarely” or “never” (Supplementary Table 3).

### Improving Reproducibility of Published Research

The majority (91%) of respondents stated that authors describing all methodology steps in detail, including any formulae analysing the data can make published science more reproducible. Around half (53%) endorsed the view that “authors should provide the source code of any custom software used to analyse the data and that the software code is well documented”, and that authors provide a link to the raw data (49%) (Supplementary figure 2). Two respondents suggested that achieving better science reproducibility would be easier if funding was more readily available for reproducing the results of others and if there were opportunities to publish the reproduced results (Supplementary section, free responses). Within the same context, some respondents recognised the current culture in science that there are not sufficient incentives in publishing reproducible (or indeed negative findings) papers, but rather being rewarded in publishing as many papers as possible in high Impact Factor journals (Supplementary section, free responses).

### Interactive Figures

Participants ranked in terms of preference potential features for an interactive figure within an article. These included choices such as “easy to manipulate” as the most preferred, and have “easy to define parameters” (Fig. 3). Generally, the answers from both the *eLIFE* and NBI surveys followed similar trends. Furthermore, free-text responses were collected, and most respondents stated that mechanisms to allow them to better understand the data presented in the figure would be beneficial, e.g. by zooming in on data (Supplementary section, free responses).

**Figure 3.**
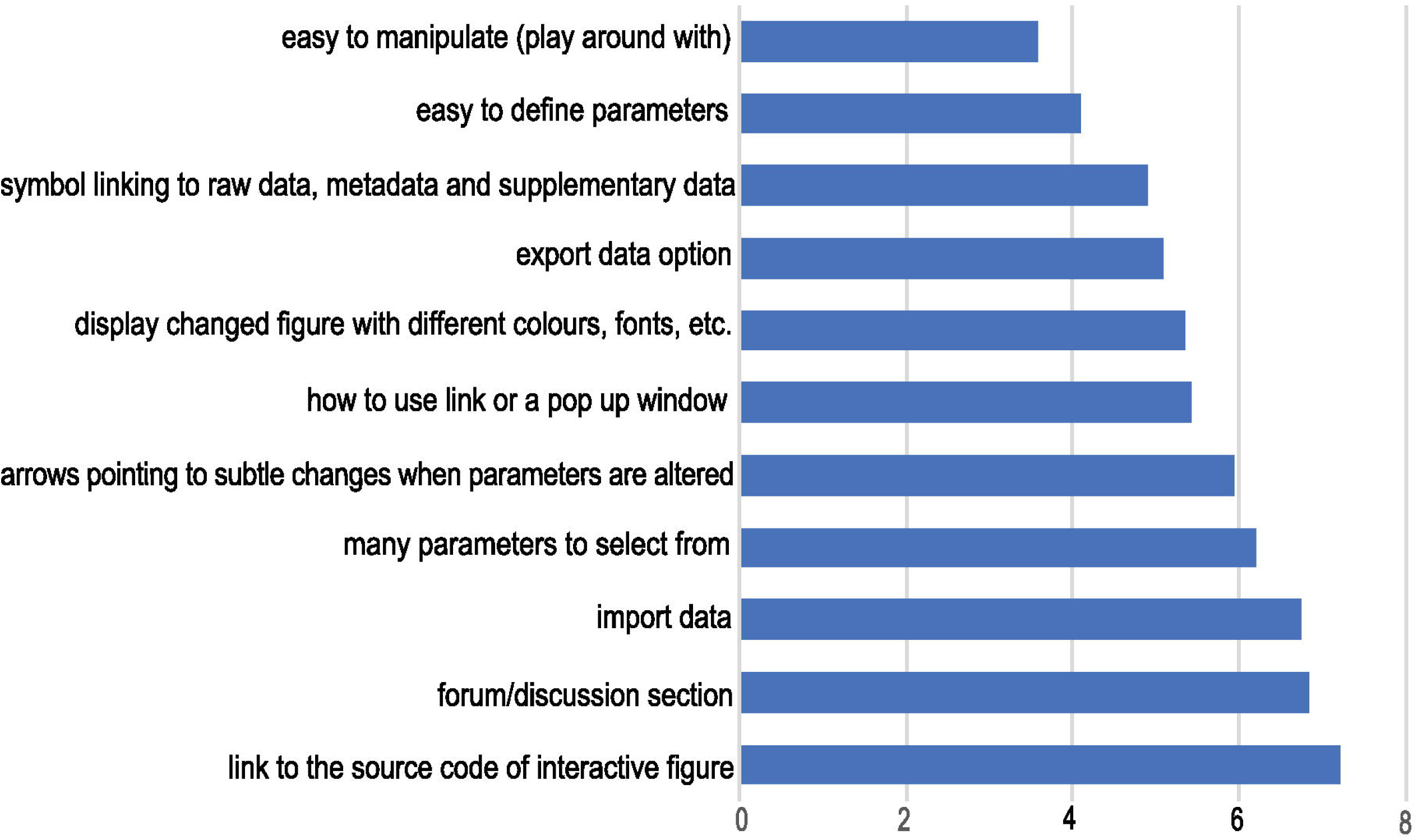
Preferred features for the interactive figure. Responses to question 9: Respondents were asked to rank in order of preference the above features, with 1 most preferred feature, to 11 the least preferred feature. The average score for each feature was calculated in order of preference as selected by the respondents from both NBI and *eLIFE* surveys. The lower the average score value (x-axis), the more preferred the feature (y-axis).

The majority of the respondents perceive a benefit in having interactive figures in published papers for both readers and authors (Fig. 4). Examples of insights included: the interactive figure would allow visualising further points on the plot from data in the supplementary section, as well as be able to alter the data that is presented in the figure; having an interactive figure like a movie or to display protein 3D structures, would be beneficial to readers. The remaining responses we categorised as software related, which included suggestions of software that could be used to produce a figure that can be interactive, such as R Shiny (shiny.studio.com). We received a total of 114 free-text responses about the respondents’ opinions on what interactive figures are and a proportion of those (25%) suggested that they had never seen or interacted with such a figure before, and no indication was given that an interactive figure would help their work (see Supplementary section, free responses).

**Figure 4.**
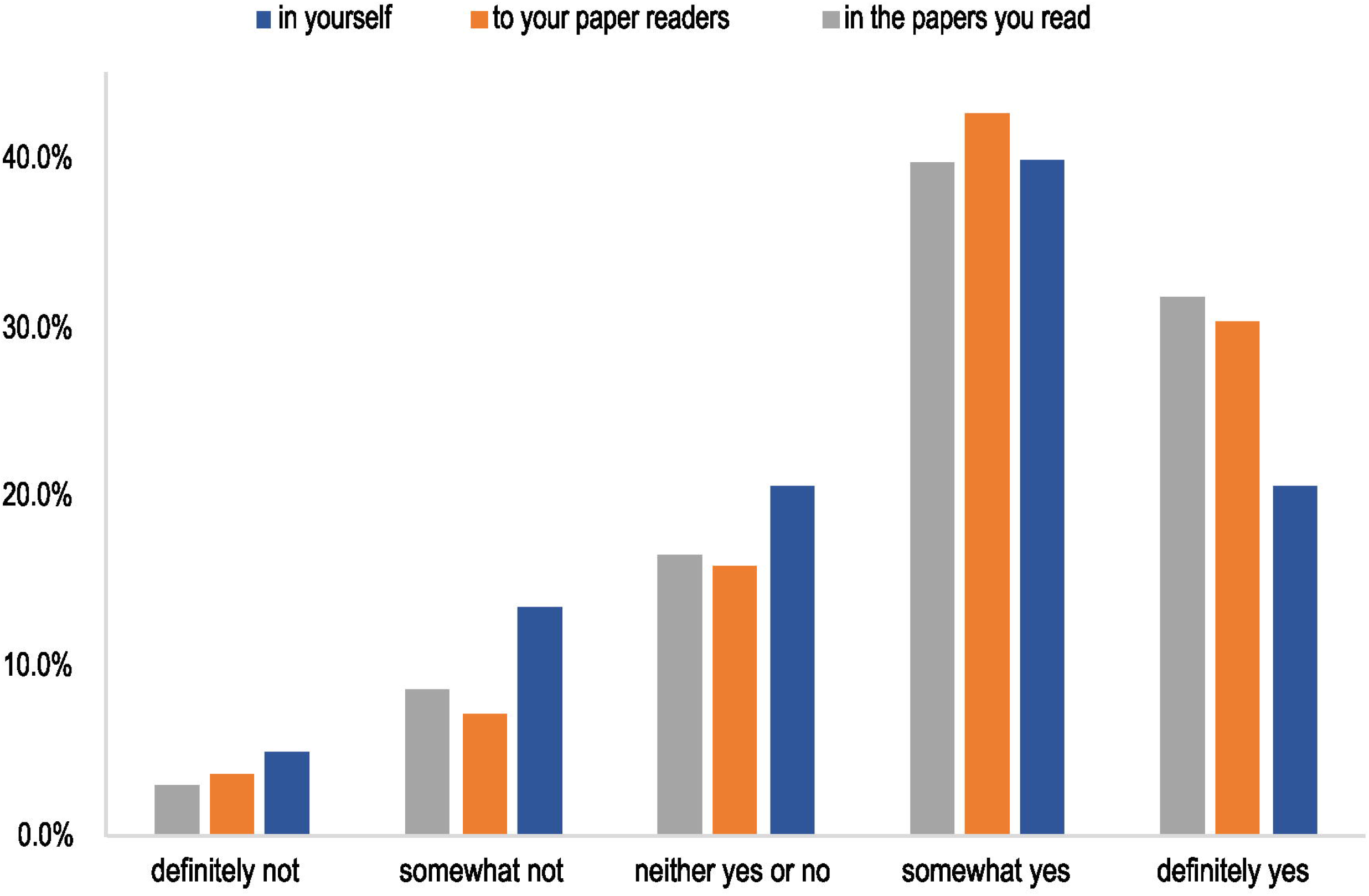
The level of perception of benefit to having the ability to publish papers with interactive figures. The benefit to the author, to the readers of the author’s papers and to the papers the author reads. Answers include the responses from both NBI and *eLIFE* surveys for question 11.

The majority of the respondents also said that they see a benefit in automatically reproducing computational experiments, and manipulating and interacting with parameters in computational analysis workflows. Equally favourable was to be able to computationally reproduce statistical analyses (Fig. 5). Despite this perceived benefit, most respondents (61%) indicated that the ability to include an interactive figure would not affect their choice of a journal when seeking to publish their research.

**Figure 5.**
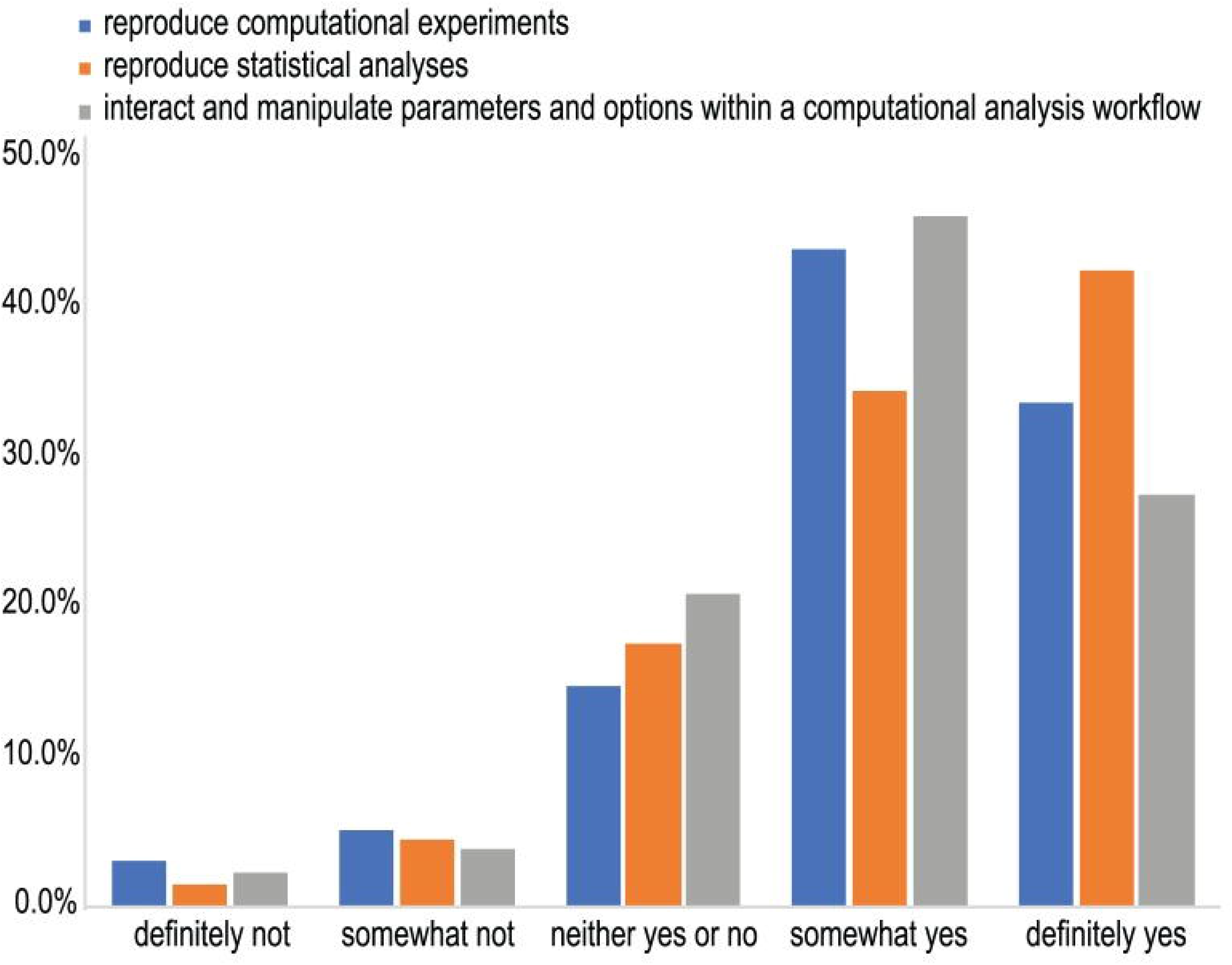
Assessment of perceived benefit for automatically reproducing computational experiments or other analyses (including statistical tests). Responses from both NBI and *eLIFE* for question 14.

### Limitations

The findings were collected using the self-reporting method which can be limited in certain ways, especially with regards to the reported reproducibility success or lack of success of the respondents. We do not know categorically that someone reproduced experiments successfully because they checked the box. Despite the potential for confusing the exact meaning of reproducibility, which could affect the answers to questions 5, 6 and 7, the general consensus among respondents showed that the questions were sufficiently phrased to help us divide people into two groups of assessment (successful vs not successful) for subsequent analysis.

Part of our survey sample were researchers from the NBI, and this population might not be representative of the life sciences research community as a whole. Researchers working in academic institutions may have attitudes, incentives or infrastructure to support reproducibility that may be different from those who work in the private sector or government agencies. In addition, as the population of the NBI researchers was solely UK based, the attitudes of these researchers might differ from those in the rest of the world, despite the fact that the NBI comprises scientists who are from multiple countries and have trained and worked in global institutions. *eLIFE* authors work across the breadth of scientific institutions, both private and public, from the international stage, thus we believe that both *eLIFE* and NBI participants to be sufficiently representative for the purposes of our survey study.

Although we do not have distinct evidence that the *eLIFE* authors cohort had a predisposition to reproducibility, and the authors we surveyed were randomly selected, we acknowledge that as *eLIFE* is a journal that requires data sharing and is also heavily involved in reproducibility efforts, such as the Reproducibility Project: Cancer Biology. In the absence of data to the contrary, thus it is reasonable to assume that some factors might have influenced the *eLIFE* respondents’ opinions about reproducibility. We do not think that this fact undermines our conclusion, but it is a factor that future studies should be aware of when drawing comparisons that can shed further light onto this issue.

We acknowledge that questions 8 and 9 were on the same page when the participants were taking the survey, and seeing the two questions together might have introduced bias into their answers. Nonetheless, free text answers to question 8 included answers which were not presented as options for question 9. Some respondents also declared that they were not aware of, or have not previously encountered, interactive figures (see Supplementary Section free text responses to question 8).

We have found that the response rate for studies of this nature is fairly typical and indeed, other studies [71-75] have experienced comparable or lower rates. Ideally, we would want to aim for a higher response rate for future studies, which could be achieved by providing monetary incentives, as well as sending email reminders to the same or bigger cohort of invited people to participate in the study [76-78].

## Discussion

This study highlights the difficulties currently experienced in reproducing experiments and conveys positive attitudes of scientists towards enabling and promoting reproducibility of published experiments through interactive elements in online publications.The NBI cohort of respondents were active life sciences researchers at the time the survey was conducted, and the *eLIFE* cohort were researchers that have published at least once in the *eLIFE* journal, therefore we believe the opinions collected are representative of researchers in life sciences who are routinely reading and publishing research.

While progress has been made in publishing standards across all life science disciplines, the opinions of the respondents reflect previously published shortcomings of the publishing procedures [79-81]: lack of data and code provision; storage standards; not including or requiring a detailed description of the methods and code structure in the published papers. However, the level of interest and incentives in reproducing published research is in its infancy, or it is not the researchers’ priority [82,83]. A key outcome of our survey is the acknowledgement of the large majority who understand that science becomes implicitly more reproducible if methods (including data, analysis, and code) are well-described and available. Respondents also perceive the benefit of having tools that enable the availability of data, methods and code and being able to automatically reproduce computational experiments described within the paper. Interactive figures within publications and executable documents can be such tools that allow the automatic reproducibility of computational experiments, or other analyses described within the paper, interact and manipulate parameters within the computational analysis workflow and give further insights and detail view of the data in the figure. Despite technologies existing to aid reproducibility and authors knowing they are beneficial, many scientific publications do not meet basic standards of reproducibility.

Our findings are in accordance with the current literature [61,84] that highlight that the lack of data access at the publication stage is one of the major reasons leading to the irreproducibility of published studies. When data is difficult to obtain, the reproducibility problem is exacerbated. A study that examined the differences between clinical and non-clinical scientists, showed that the majority of respondents did not have experience with uploading biomedical data to a repository, stemming from different social reasons not to do so: concerns and motivation around data sharing; work necessary to prepare the data [73]. Even with current policies mandating data openness [59,60], authors still fail to include their data alongside their publication, and this can not only be attributed to technical complications, but also fear of being scooped, fear of mistakes being found in data or analyses, and fear of others using their data for their own research papers [64,73]. Making FAIR data practices standard, through public data deposition and subsequent publication and citation, could encourage individual researchers and communities to share and reuse data considering their individual requirements and needs [68]. Data accessibility issues are also compounded by data becoming less retrievable with every year passing after the publication [85]. This is supported by our findings where data is either not available upon publication (57%) or authors cannot be reached/are unresponsive to data provision requests (44%). This continues to be a cultural artefact of using a paper’s methods section as a description of steps to reproduce analysis, rather than a fully reproducible solution involving easy access to public data repositories, open source code, and comprehensive documentation.

As evidenced by the respondents, the lack of data availability is a common hurdle for researchers to encounter that prevents the reproducibility of published work. Thus, the reproducibility of experiments could be improved by increasing the availability of data. Datasets are becoming larger and more complex, especially in the area of genomics. Storage solutions for large data files and citing them within the publication document, especially those in the order of terabytes, can allow for their wider, more efficient and proper data reusability [86,87]. Despite the potential advantage, these services can provide for data availability and accessibility, they do not implicitly solve the problem of data *reusability*. This is most apparent when data is too large to be stored locally or transferred via slow internet connections, or there is no route to attach metadata that describes the datasets sufficiently for reuse or integration with other datasets. There is also the question of data repository *longevity* - who funds the repositories for decades into the future? Currently, some researchers now have to pay data egress charges for downloading data from cloud providers [88-90]. This method presumably saves the data producers money in terms of storing large datasets publicly, but the cost is somewhat now presented to the consumer. This raises complex questions around large data generation projects that also need to be studied extensively for future impact, especially with respect to reproducibility within publications. Moreover, access to the raw data might not be enough, if the steps and other artefacts involved in producing the processed data that was used in the analysis are not provided [68]. In addition, corresponding authors often move on from projects and institutions or the authors themselves can no longer access the data, meaning “data available on request” ceases to be a viable option to source data or explanations of methods. Restricted access to an article can also affect reproducibility by requiring paid subscriptions to read content from a publisher. Although there is precedent for requesting single articles within cross-library loan systems or contacting the corresponding author(s) directly, this, much like requesting access to data, is not without issues. Pre-print servers such as bioRxiv have been taken up rapidly [91], especially in the genomics and bioinformatics domains, and this has the potential to remove delays in publication whilst simultaneously providing a “line in the sand” with a Digital Object Identifier (DOI) and maintaining the requirements for FAIR data. In some cases, the sensitivity of data might discourage authors from data sharing [92,93], but this reason was only reported by a small proportion of our respondents. Whilst there are efforts that attempt to apply the FAIR principles to clinical data, such as in the case of the OpenTrials database [94], they are by no means ubiquitous.

Data within public repositories with specific deposition requirements (such as the EMBL-EBI European Nucleotide Archive, ebi.ac.uk/ena), might not be associated or annotated with standardised metadata that describes it accurately [95], rather the bare minimum for deposition. Training scientists to implement data management policies effectively is likely to increase data reuse through improved metadata. In a 2016 survey of 3987 National Science Foundation Directorate of Biological Sciences principal investigators (BIO PIs), expressed their greatest unmet training needs by their institutions [83]. These were in the areas of integration of multiple data (89%), data management and metadata (78%) and scaling analysis to cloud/high-performance computing (71%). The aforementioned data and computing elements are integral to the correct knowledge “how-to” for research reproducibility. Our findings indicated that those who stated they had experience in informatics also stated they are better able to attempt and reproduce results. Practical bioinformatics and data management training, rather than in specific tools, may be an effective way of reinforcing the notion that researchers’ contributions towards reproducibility are a responsibility that requires active planning and execution. This may be especially effective when considering the training requirements of wet-lab and field scientists, who are becoming increasingly responsible for larger and more complex computational datasets. Further research needs to be undertaken to better understand how researchers’ competence in computational reproducibility may be linked to their level of informatics training.

Furthermore, there remains a perception that researchers do not get credit for reproducing the work of others or publishing negative or null results [96,97]. Whilst some journals explicitly state that they welcome negative results articles (e.g. PLOS One “Missing Pieces” collection), this is by no means the norm in life science publishing as evidenced by low, and dropping publication rates of negative findings [96-98]. In addition, the perception that mostly positive results are publication-worthy might discourage researchers from providing enough details on their research methodology, such as reporting any negative findings. For transparent and reproducible science, both negative (or null) and positive results should be reported for others to examine the evidence [96,99,100]. Ideally, the publication system would enable checking of reproducibility by reviewers and editors at the peer-review stage, with authors providing all data (including raw data), a full description of methods including statistical analysis parameters, any negative findings based on previous work and open source software code [101]. These elements can all be included within the interactive figure, such as by zooming in on over data points to reveal more information on the data, pop up windows to give details on negative results and parameters and the figure offering re-running of the computational experiment in the case of executable documents. Peer reviewers would then be better able to check for anomalies, and editors could perform the final check to ensure that the science paper to be published is presenting true, valid, and reproducible research. Some respondents have suggested that if reviewers and/or editors were monetarily compensated, spending time to reproduce the computational experiments in manuscripts would become more feasible, and would aid the irreproducibility issue. However, paying reviewers does not necessarily ensure that they would be more diligent in checking or trying to reproduce results [102] and there must be optimal ways to ensure effective pressure is placed upon the authors and publishing journals to have better publication standards [103,104]. The increasing adoption by journals of reporting standards for experimental design and results, provide a framework for harmonising the description of scientific processes to enable reproducibility. However, these standards are not universally enforced [105]. Similarly, concrete funding within research grants for implementing reproducibility itself manifested as actionable Data Management Plans (dcc.ac.uk, 2019), rather than what is currently a by-product of the publishing process, could give a level of confidence to researchers who would want to reproduce previous work and incorporate that data in their own projects.

Respondents mentioned that there are word count restrictions in papers, and journals often ask authors to shorten methods sections and perhaps move some text to supplementary information, many times placed in an unorganised fashion or having to remove it altogether. This is a legacy product of the hard-copy publishing era and readability aside, word limits are not consequential for most internet journals. Even so, if the word count limit was only applicable to the introduction, results and discussion sections, then the authors could describe methods in more detail within the paper, without having to move that valuable information in the supplementary section. When methods are citing methodology techniques as described in other papers, where those original references are hard to obtain, typically through closed access practices or by request mechanisms as noted above, then this can be an additional barrier to the reproducibility of the experiment. This suggests that there are benefits to describing the methods in detail and stating that they are similar to certain (cited) references as well as document the laboratory’s expertise in a particular method [106]. However, multi-institutional or consortium papers are becoming more common with ever-increasing numbers of authors on papers, which adds complexity to how authors should describe every previous method available that underpins their research [100]. There is no obvious solution to this issue. Highly specialised methods (e.g. electrophysiology expertise, requirements for large computational resources or knowledge of complex bioinformatics algorithms) and specific reagents (e.g. different animal strains), might not be readily available to other research groups [83]. As stated by some respondents, in certain cases the effective reproducibility of experiments is obstructed by numerical issues with very small or very large matrices or datasets, or different versions of analysis software used, perhaps to address bugs in analytical code, will cause a variation in the reproduced results.

Previous studies have provided strong evidence that there is a need for better technical systems and platforms to enable and promote the reproducibility of experiments. We provide additional evidence that that paper authors and readers perceive a benefit from having an interactive figure that would allow for the reproducibility of the experiment shown in the figure. An article that gives access to the data, code and detailed data analysis steps would allow for *in situ* reproduction of computational experiments by re-running code including statistical analyses “live” within the paper [26]. Whilst our study did not concentrate on how these “executable papers” may be constructed, this is an active area of development and some examples of how this may be achieved have been provided [51,108]. We provide additional evidence that paper authors and readers perceive a benefit from having publication infrastructure available that would allow for the reproducibility of an experiment. As such, the findings of this survey helped the development of two prototypes of interactive figures (see Data and Code availability) and subsequently the creation of *eLIFE*’s first computationally reproducible document [52].

We also asked whether presenting published experiments through interactive figures elements in online publications might be beneficial to researchers, in order to better consume research outputs. Respondents stated they could see the benefit in having interactive figures for the readers of their papers and the papers they read and being able as authors to present their experiment analysis and data as interactive figures. Respondents endorsed articles which include interactive elements, where access to the processed and raw data, metadata, code, and detailed analysis steps, in the form of an interactive figure, would help article readers better understand the paper and the experimental design and methodology. This would, in turn, improve the reproducibility of the experiment presented in the interactive figure, especially computational experiments. The notion of data visualisation tools promoting interactivity and reproducibility in online publishing has also been discussed in the literature [42]. Other efforts have been exploring the availability of interactive figures for driving reproducibility in publishing in the form of executable documents [28,51,52,55]. Moreover, technologies such as Jupyter Notebooks, Binder, myExperiment, CodeOcean enable the reproducibility of computational experiments associated with publications, provided by the authors as links from the paper. However, the benefit of having the interactivity and availability of reproducing experiments from within the article itself in the form of interactive figures, is that the reader can stay within the article itself and explore all the details of the data presented in the figure, download the data, play with the code or analysis that produced the figure, interact with parameters in the computational analysis workflows and computationally reproduce the experiments presented in the figure. This can enable the reader to better understand the research done presented in the interactive figure. Despite the self-reported perceived benefits of including interactive figures, the availability of this facility would not affect the respondents’ decisions on where to publish. This contradiction suggests that cultural factors (incentives, concerns authors have with sharing their data, attitudes towards open research) [64,73] play an underestimated role in reproducibility.

Despite the benefits, the interactive documents and figures can provide to the publishing system for improved consumption of research outputs, and that those benefits are in demand by the scientific community, work is needed in order to promote and support their use. Given the diversity of biological datasets and ever-evolving methods for data generation and analysis, it is unlikely that a single interactive infrastructure type can support all types of data and analysis. More research into how different types of data can be supported and presented in papers with interactivity needs to be undertaken. Yet problems with data availability and data sizes will persist - many studies comprise datasets that are too large to upload and render within web browsers in a reasonable timescale. Even if the data are available through well-funded repositories with fast data transfers, e.g. the INSDC databases (insdc.org), are publishers ready to bear the extra costs of supporting the infrastructure and people required to develop or maintain such interactive systems in the long run? These are questions that need to be further investigated, particularly when considering any form of industry standardisation of such interactivity in the publishing system. It is clear that publishing online journal papers with embedded interactive figures requires alterations to infrastructure, authoring tools and editorial processes [42]. In some cases, the data underpinning the figures might need to be stored and managed by third parties and this means the data, as well as the figures, may not be persistent. The same argument is relevant to software availability and reuse - publishers would need to verify that any links to data and software were available and contained original unmodified datasets. As datasets become larger and more complex, and more software and infrastructure is needed to re-analyse published datasets, this will affect how infrastructure will need to be developed to underpin reproducible research. Incentives will need to be put in place to motivate investment in these efforts.

We show that providing tools to scientists who are not computationally aware also requires a change in research culture, as many aspects of computational reproducibility require a change in publishing behaviour and competence in the informatics domain. Encouraging and incentivising scientists to conduct robust, transparent, reproducible and replicable research, such as with badges to recognise open practices should be prioritised to help solve the irreproducibility issue [109]. Implementing hiring practices with open science at the core of research roles [110] will encourage attitudes to change across faculty departments and institutions.

Another potential solution to the reproducibility crisis is to identify quantifiable metrics of research reproducibility and its scientific impact, thus giving researchers a better understanding of how their work stands on a scale of measurable reproducibility. The current assessment of the impact of research articles is a set of quantifiable metrics that do not evaluate research reproducibility, but stakeholders are starting to request that checklists and tools are provided to improve these assessments [111]. It is harder to find a better approach that is based on a thoroughly informed analysis by unbiased experts in the field that would quantify the reproducibility level of the research article [112]. That said, top-down requirements from journals and funders to release reproducible data and code may go some way to improving computational reproducibility within the life sciences, but this will also rely on the availability of technical solutions that are accessible and useful to the majority of scientists.

Opinions are mixed regarding the extent and severity of the reproducibility crisis. Our study and previous studies are highlighting the need to find effective solutions towards solving the reproducibility issue. Steps towards modernising the publishing system by incorporating interactivity with interactive figures and by automatically reproducing computational experiments described within a paper are deemed desirable. This may be a good starting point for improving research reproducibility by reproducing experiments within research articles. This, however, does not come without its caveats, as we described above. From our findings, and given the ongoing release of tools and platforms for technical reproducibility, future efforts should be spent in tackling the cultural behaviour of scientists, especially when faced with the need to publish for career progression.

## Acknowledgements

This project is funded by a BBSRC iCASE Studentship (project reference: BB/M017176/1). We would like to thank all the respondents of the surveys for their time. We would also like to thank George Savva from the Quadram Institute (QIB, UK) for comments and suggestions for this manuscript; *eLIFE* Sciences Publications Ltd, with whom the corresponding author collaborates as an iCASE student; as well as Ian Mulvany, former *eLIFE* Head of Development, for his help in developing the survey questionnaire.

## Data and Code Availability

All data files are available via this Url: https://doi.org/10.6084/m9.figshare.c.4436912. Prototypes of interactives figures developed by the corresponding author are available via these GitHub repositories: https://github.com/code56/nodeServerSimpleFig and https://github.com/code56/prototype_article_interactive_figure

